# The connection between Rap1 and Talin1 in CD4^+^ T Lymphocytes

**DOI:** 10.1101/2021.09.22.461411

**Authors:** Frederic Lagarrigue, Boyang Tan, Qinyi Du, Zhichao Fan, Miguel A. Lopez-Ramirez, Alexandre R Gingras, Weiwei Qi, Hao Sun

**Author notes:** Correspondence: Hao Sun.

## Abstract

Agonist induced increase in integrin affinity for ligands (activation) plays a pivotal role in T cell trafficking and functions. Activation requires Rap1 GTPase-mediated recruitment of talin1 to the integrins in the plasma membrane. Rap1-interacting adaptor molecule (RIAM) is a Rap1 effector that serves this function in T cells. In addition, Rap1 directly binds to talin1 to enable integrin activation in platelets. Here, we assessed the relative contributions of the Rap1-talin1 interaction and RIAM and provide a complete accounting of the connections between Rap1 and talin1 that support integrin activation in conventional CD4^+^ (Tconv) and CD25^Hi^Foxp3^+^CD4^+^ regulatory T (Treg) cells. Disruption of both Rap1 binding sites in talin1 (talin1 (R35E,R118E)) causes a partial defect in αLβ2, α4β1 and α4β7 integrin activation in both Tconv and Treg cells with resulting defects in T cell homing and functions. Over-expression of RIAM bypasses the integrin activation defect in Tconv cells expressing talin1 (R35E,R118E), indicating that RIAM can substitute for Rap1 binding to talin in integrin activation. Conversely, deletion of RIAM in talin1 (R35E,R118E) Tconv cells abrogates activation of αLβ2, α4β1 and α4β7. RIAM and lamellipodin (LPD) are mammalian members of the MRL protein family; LPD plays a more important role than RIAM in Treg cell integrin activation. Nevertheless, loss of RIAM profoundly exacerbates the defects in Treg cell function caused by the talin1 (R35E,R118E) mutation. Most importantly, deleting both MRL proteins combined with talin1 (R35E,R118E) phenocopies the complete lack of integrin activation observed in Rap1a/b null Treg cells. In sum, these data reveal the functionally significant connections between Rap1 and talin1 that enable αLβ2, α4β1 and α4β7 integrin activation in T cells.

## Introduction

Integrin adhesion receptors play an essential role in lymphocytes to shape a successful adaptative immune response (1, 2). Integrins participate in lymphocyte development and trafficking to lymphoid organs and sites of inflammation (3, 4). In addition, they are instrumental in the formation of immunological synapses to control T cell functions (5), in particular their killing activity and capacity to present antigens. Binding of αLβ2 (LFA-1, CD11a/CD18), α4β1 (VLA-4) and α4β7 integrins to their cognate ligands is precisely controlled in lymphocytes (6, 7), and deregulation in their activity contributes to various diseases such as multiple sclerosis, asthma, atherosclerosis and inflammatory bowel disease (8-10).

Integrins in lymphocytes are usually maintained in a low-affinity state until agonist stimulation induces a high-affinity form (11). This process operationally defined as “integrin activation” is central to T cell functions (6, 7). The capacity of intracellular signaling pathways to induce such changes in integrin conformation and affinity depends on binding of the cytosolic adapter talin1 to the integrin β cytoplasmic tail, which is a critical common final step in integrin activation (12, 13). Talin1 is unique in its ability to regulate activation of β1 (13), β2 (14, 15), β3 (13), and β7 (16) integrins. Talin1 is a large multi-domain protein that links integrins to the actin cytoskeleton via its N-terminal head domain that binds to the integrin β cytoplasmic tail and its C-terminal rod domain that binds F-actin (17, 18). Talin1 head domain (THD) comprises an atypical FERM domain characterized by an additional F0 subdomain to the characteristic FERM F1, F2 and F3 subdomains (17). Talin1 is autoinhibited in the cytosol due to the interaction of the talin1 head domain (THD) with the rod domain, which prevents its interaction with the integrin β cytoplasmic tail (19). The effects of talin1 deletion in mouse blood cells have unequivocally confirmed its importance in integrin activation in lymphocytes (14, 20-22); however, the sequence of signal transduction events from adhesive stimuli to talin1 recruitment to integrins are not yet fully characterized.

T cell stimulation via chemokines or antigen receptors initiates a cascade of signaling pathways that converge on talin1 to trigger integrin activation (2). Rap1 GTPases function as a prominent signaling hub that controls talin1 binding to T cell integrins (23). Pioneer studies identified Rap1-interacting adaptor molecule (RIAM) as a Rap1 effector that recruits talin1 to the plasma membrane to enable its interaction with the integrin β cytoplasmic tail (24). RIAM is abundant in hematopoietic cells (25). In contrast to talin1 deletion (26, 27), germline loss of RIAM in mice does not affect development, hemostasis or platelet integrin activation and function (28-30). However, RIAM null mice exhibit a significant leukocytosis and impaired integrin-mediated T cell adhesion and homing to lymph nodes (29, 30). Moreover, RIAM plays an essential role in the dynamic regulation of lymphocyte integrin function during the formation of the immunological synapse and killing activity (31). RIAM deficiency results in a loss of β2 integrin activation in multiple leukocyte subsets including T cells, whereas activation of α4β1 integrin operates in a RIAM-independent manner (29, 30). In contrast, we showed that RIAM is dispensable for integrin activation and function in regulatory T (Treg) cells despite the fact that Rap1 is required for Treg cell function and RIAM is expressed in Treg cells (32). Increased expression of Lamellipodin (LPD), a RIAM paralogue, accounts for the lack of RIAM requirement in Treg cells (32). Accordingly, targeting RIAM could provide an avenue for cell-type specific inhibition of integrin activation to ameliorate auto-immune disorders such as type I experimental diabetes (31) or inflammatory bowel disease (32) by suppressing integrin-mediated homing and function of conventional T (Tconv) cells while sparing Treg cells.

Because RIAM-deficiency in T cells has a less pronounced effect on leukocyte β2 integrin function than loss of either talin1 or Rap1 expression (28, 29), the existence of alternative RIAM-independent pathways that regulate Rap1-mediated recruitment of talin1 to T cell integrins becomes evident. Previous studies revealed that Rap1 can bind directly to talin1 F0 subdomain with low affinity (33, 34). However, blockade of Rap1-talin1 F0 interaction has a relatively minor effect on platelet integrin activation and hemostasis (35, 36). Thereafter, we identified a second Rap1 binding site in talin1 F1 subdomain of similar affinity to that in F0 (37). Mutation R118E in talin1 F1 subdomain, which blocks Rap1 binding, abolished the capacity of Rap1 to mediate talin1-induced integrin activation in model CHO cells and platelets (37, 38). Disruption of Rap1 binding to both talin1 F0 and F1 subdomains further impairs integrin activation to a similar extent to that observed in platelets lacking both Rap1a and Rap1b (38). These studies revealed that Rap1 binding to both talin1 F0 and F1 subdomains is crucial for platelet integrin function and hemostasis. In addition to likely facilitating membrane recruitment of talin1 in platelets and endothelial cells, the direct interaction of talin1 with Rap1 can place talin1 in proximity to and in appropriate orientation with plasma membrane phospholipids and the integrin β3 cytoplasmic tail, leading to integrin activation (39).

Here, we assessed the contribution of the Rap1-talin1 interaction to trigger integrin activation in T cells. We generated mice bearing point mutations in talin1 F0 subdomain (R35E), F1 subdomain (R118E) or both F0 and F1 subdomains (R35E,R118E) (35, 38). Disruption of Rap1 binding to either talin1 F0 or F1 subdomains causes a significant, but moderate, defect in αLβ2, α4β1 and α4β7 integrin activation in CD4^+^ T cells. T cells expressing the talin1(R35E,R118E) mutant manifest further impaired integrin activation. Overexpression of RIAM in mutant T cells expressing talin1(R35E,R118E) restored the capacity of Rap1 to potentiate integrin activation. In contrast, loss of RIAM in talin1(R35E,R118E) mutant T cells exacerbate the defects in integrin activation. Thus, direct binding of Rap1 to talin1 operates as an alternative pathway to the Rap1-RIAM axis to control integrin activation in CD4^+^ T cells. Similarly, Rap1-talin1 interaction participates, in parallel with LPD, to αLβ2 and α4β7 integrin function in Treg cells. These data reveal the contribution of multiple routes that connect Rap1 to talin1 in Treg and CD4^+^ T cells and highlight the potential of interfering with these pathways to specifically manipulate the trafficking and function of selective lymphocyte subsets to ameliorate autoimmunity.

## Materials and Methods

### Antibodies and reagents

The following antibodies were from BioLegend: CD3 (17A2, 2C11), CD4 (GK1.5), CD29 (HMb1-1), CD49d (R1-2), β7 integrin (FIB504) and Foxp3 (MF-14). Antibody against RIAM was from R&D Systems. Antibodies against talin1 (8d4, Sigma-Aldrich), RIAM (produced in rabbits immunized with recombinant 1-301 portion of human RIAM), Rap1 (sc-398755, Santa Cruz) and β-Actin (A5316, Sigma) were used for Western blot. Secondary antibodies conjugated to Alexa Fluor were from Jackson ImmunoResearch. eFluor 670 was from Biolegend. PMA was from Sigma-Aldrich. MojoSort mouse CD4 T cell isolation kit was from BioLegend. Liberase TL (Research Grade) and DNAseI were from Roche. Recombinant mouse VCAM-1-Fc was from R&D Systems. Recombinant mouse ICAM-1 and VCAM-1-Fc were from BioLegend. Recombinant mouse MAdCAM-1 was from R&D Systems.

### Mice

All animal experiments were approved by the Institutional Animal Care and Use Committee (IACUC) of the University of California, San Diego, and were conducted in compliance with federal regulations as well as institutional guidelines and regulations on animal studies. *Apbb1ip*^*Flox/Flox*^ (28, 29), *Tln1*^*Flox/Flox*^ (26), *Tln1*^*wt/R35E*^ (35), *Tln1*^*wt/R118E*^ (38), *Tln1*^*wt/R35E,R118E*^ (38), *Rap1a/b*^*Flox/Flox*^ (29), *Tln1*^*wt/L325R*^ (40), *Raph1*^*Flox/Flox*^ (41), *CD4*^*Cre*^ (42) and *Foxp3*^*YFP-Cre*^ (43) mice have been described previously. Hemizygous mice with point mutations in Tln1 were crossed with *Tln1*^*Flox/Flox*^; *CD4*^*Cre*^ to obtain lymphocyte-specific deletion of Tln1 flox allele in *Tln1*^*Flox/wt*^; *CD4*^*Cre*^ (control) and *Tln1*^*Flox/Mut*^; *CD4*^*Cre*^ (Tln1-cMut) littermates for experiments. Similarly, *Tln1*^*Flox/wt*^; *Foxp3*^*YFP-Cre*^ (control) were compared to *Tln1*^*Flox/Mut*^; *Foxp3*^*YFP-Cre*^ (Tln1-rMut) littermates for experiments. RIAM-cKO refers to *Apbb1ip*^*Flox/Flox*^; *CD4*^*Cre*^ mice, and RIAM-rKO refers to *Apbb1ip*^*Flox/Flox*^; *Foxp3*^*YFP-Cre*^. Similarly, LPD-rKO refers to *Raph1*^*Flox/Flox*^; *Foxp3*^*YFP-Cre*^. Eight-twelve-week-old sex-matched wild-type littermates were control animals for all experiments. All mice were housed in specific pathogen-free conditions.

### Blood counts

Peripheral blood was collected from the retro-orbital plexus into tubes containing K^+^/EDTA. Cell counts were performed using a Hemavet 950FS Hematology System programmed with mouse-specific settings (Drew Scientific). All samples were tested in duplicate, and the mean for each animal was plotted.

### T cell purification

T cells were isolated from spleens as previously described (32). Briefly, CD4^+^ T cells were purified by magnetic separation using the CD4^+^ T cell negative isolation kit (Biolegend) and were routinely > 95% CD3^+^ by flow cytometry. CD4^+^CD25^-^ T cells (Tconv) were purified by adding a biotinylated anti-CD25 antibody (PC61; Biolegend) to the CD4^+^ T cell negative isolation kit to deplete Treg cells. YFP^+^ Treg cells were sorted with a FACSAria 2 (BD Biosciences).

### T cell transduction

Human RIAM cDNA was fused to EGFP at the C-terminus and cloned downstream to the murine EF1α promoter into the pMSCV retroviral backbone. 293-LVX cells (Clontech) were transfected with pMSCV and pCl-Eco plasmids using Lipofectamine 3000 (Invitrogen) and cultured for 3 days. Purified CD4^+^ T cells were activated with immobilized anti-CD3 mAb (145-2C11 clone, 5 μg/ml), anti-CD28 mAb (37.51 clone, 5 μg/ml) and IL-2 (20 U/ml) in RPMI media for 2 days. Then the cells were washed, mixed with 50% viral supernatant, 50% fresh RPMI medium supplemented with 20 U/ml IL-2 and spin-fected on RetroNectin (Takara)-coated plates at 3000 *g* at 32°C for 2 h. After 1 day, the transduction step was repeated. The next day, cells were washed and maintained in culture between 8 ×10^5^ and 1×10^6^ cells per ml with IL-2 (20 U/ml). Cells were assayed for transduction efficiency and function by flow cytometry 2-3 days after the second transduction.

### Flow cytometry

Cells isolated from mouse tissues were washed and resuspended in PBS containing 0.1% BSA and stained with conjugated antibody for 30 min at 4°C. Then, cells were washed twice before data acquisition. Intracellular staining with anti-RIAM antibody was performed in 0.5% saponin after fixation with 2% formaldehyde. For soluble ligand binding assay, 5 × 10^6^ cells were washed and resuspended in HBSS containing 0.1% BSA and 1 mM Ca^2+^/Mg^2+^ before incubation with integrin ligands (5 μg/ml) for 30 min at 37°C with or without 100 nM PMA. Cells were then incubated with Alexa Fluor 647–conjugated anti-human IgG (1:200) for 30 min at 4°C. Flow cytometric analysis was performed using an Accuri C6 Plus (BD Biosciences). Data were analyzed using FlowJo software.

### Flow chamber assay

Polystyrene Petri dishes were coated with murine VCAM-1/Fc, ICAM-1/FC or MAdCAM-1/Fc (10 μg/ml) alone or with CXCL12 (SDF1α, 1 μg/ml) in coating buffer (PBS, 10 mM NaHCO_3_, pH 9.0) for 1 h at room temperature, blocked for 1 h with BSA (1%), and then assembled into the flow chamber device (GLycoTech). Cells were diluted to 5×10^6^ cells/ml in HBSS (with 1 mM Ca^2+^/Mg^2+^) and immediately perfused through the flow chamber at a constant flow of 2 dyn/cm^2^. For the PMA stimulation, cells were pre-stimulated for 10 min with PMA (100 nM) at 37°C before perfusion. Adhesive interactions between the flowing cells and the coated substrates were assessed by manually tracking the motions of individual cells for 1 min as previously described (44). Cells that remained adherent and stationary after the initial adhesion point for >10 s with a velocity <1 μm/s were defined as arrested adherent cells.

### In vivo competitive lymphocyte homing

Mutant and WT T cells were labeled with 1 or 10 μM CFSE, respectively, resulting in readily discriminated cell populations. Equal numbers (1×10^7^) of differentially labeled cells were mixed and then intravenously injected into C57BL/6 recipient mice. Lymphoid organs were harvested 3 h after injection, and isolated cells were analyzed by flow cytometry for the ratio of mutant (CFSE^lo^) to WT (CFSE^hi^) T cells in various lymphoid organs.

### Treg suppression assay

CD4^+^CD25^-^ T cells (Responder cells) were isolated from spleens of C57BL/6 (CD45.1) WT mice. Responder cells were labelled with CFSE and cocultured with YFP^+^ Treg cells (8:1, 4:1, 2:1 and 1:1 ratios) in the presence of 5 μg/ml immobilized anti-CD3 mAb (145-2C11 clone, 5 μg/ml), anti-CD28 mAb (37.51 clone, 5 μg/ml), and IL2 (100 U/ml) for 4 days at 37°C. Proliferation index was calculated using FlowJo v10 software.

### Western blot

Isolated CD4^+^ T cells and sorted Tregs were pelleted by centrifugation at 700 g for 5 min at room temperature and then lysed in Laemmli sample buffer. Lysates were subjected to a 4-20% gradient SDS-PAGE. The appropriate IRDye/Alexa Fluor-conjugated secondary antibodies were from LI-COR. Nitrocellulose membranes were scanned using an Odyssey CLx infrared imaging system (LI-COR) and blots were processed using Image Studio Lite software (LI-COR).

### Statistical Analysis

Statistical significance was assayed by a two-tailed *t*-test for single comparisons. ANOVA with a Bonferroni post hoc test was used to assay statistical significance for multiple comparisons. All datasets were tested for Gaussian normality distribution. Statistical analysis was performed using Prism software (version 8.0, GraphPad). A *p* value <0.05 was considered significant.

## Results

### Rap1 binding to the talin-1 F0 or F1 domain contributes to integrin activation in CD4^+^ T cells

We previously generated knock-in mouse strains harboring the point mutations R35E and R118E in talin1 F0 and F1 subdomains (Fig. 1A), respectively, to disrupt binding to Rap1 (35, 38). We crossed *Tln1*^*WT/R35E*^*;CD4-Cre*^*+/-*^ mice with the *Tln1*^*Flox/Flox*^ strain to T cell-specific deletion of Tln1 flox allele in *Tln1*^*WT/Flox*^*;CD4-Cre*^*+/-*^ (control) and *Tln1*^*R35E/Flox*^*;CD4-Cre*^*+/-*^ (indicated as Tln1-cR35E) littermates for experiments. Similarly, *Tln1*^*R118E/Flox*^*;CD4-Cre*^*+/-*^ mice (indicated as Tln1-cR118E) were compared to *Tln1*^*WT/Flox*^*;CD4-Cre*^*+/-*^ (control) littermates for experiments. *Tln1*^*WT/R35E*^*;CD4-Cre*^*+/-*^ and *Tln1*^*R118E/Flox*^*;CD4-Cre*^*+/-*^ progenies selectively express talin1(R35E) or talin1(R118E) mutant proteins in T cells. These mutant mice were viable, exhibited no gross developmental defects and appeared healthy. Lymphocyte expression of talin1, Rap1a/b and RIAM was unchanged (Supplementary Fig. 1A). Surface expression of αL, β2, α4, β1 and β7 integrins in Tln1-cR35E and Tln1-cR118E mice was similar to control T cells (Supplementary Fig. 1B). These data indicate that both Tln1-cR35E and Tln1-cR118E mice are suitable for examination of the effects of blocking Rap1 binding to talin1 F0 and F1 subdomains on T cell integrin activation and function.

**Figure 1.**
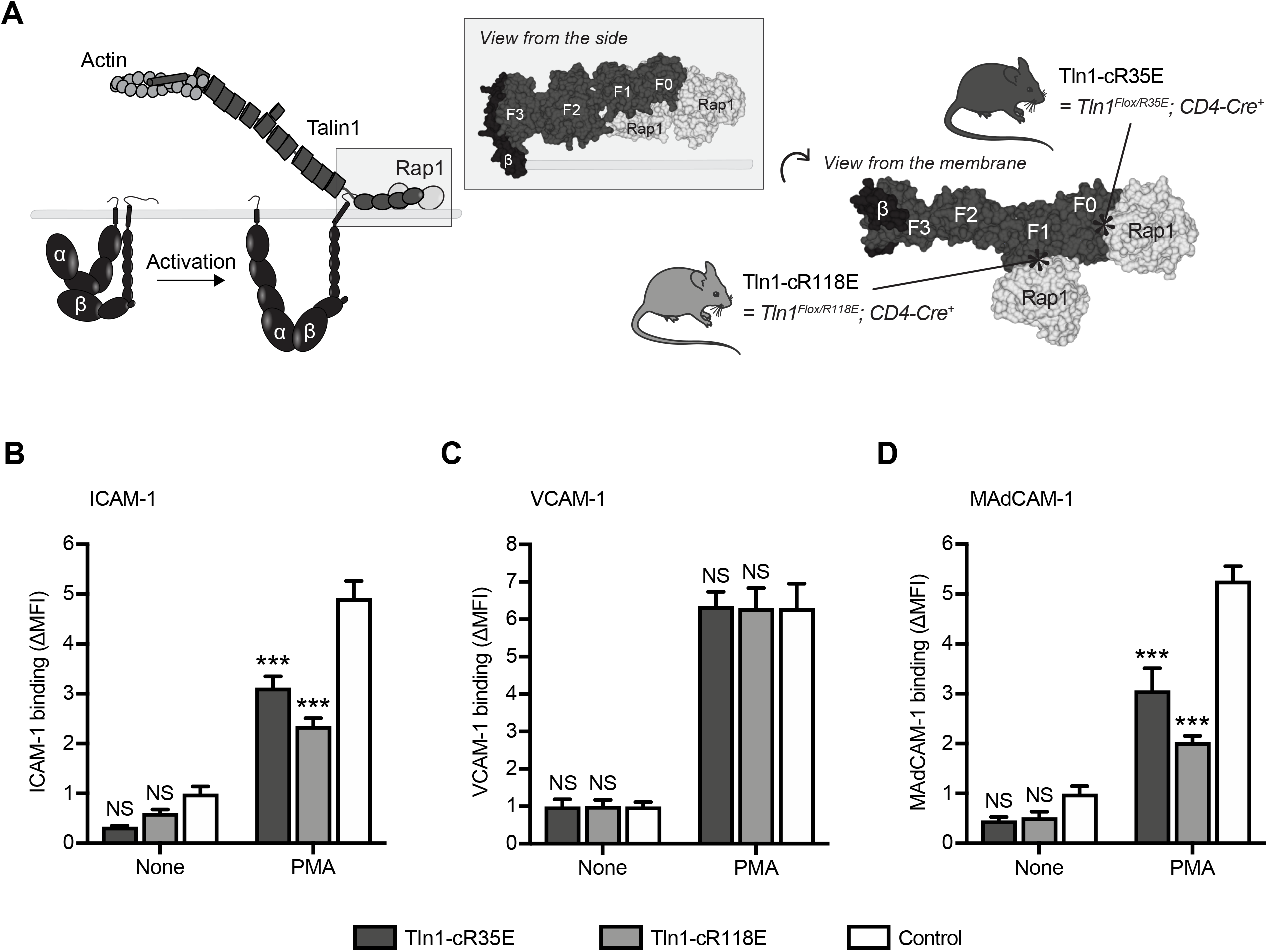
Rap1 binding to talin1 F0 or F1 domain contributes to integrin activation in CD4^+^ T cells and Treg cells. (A) Diagram depicting the role of talin1 in integrin activation. Mutations R35E and R118E in Talin1 block Rap1 binding to F0 and F1 domains, respectively. (B-D) Binding of control and Tln1-cR35E or Tln1-cR118E mutant CD4^+^ T cells to soluble integrin ligands. Cellular stimulation was achieved with 100 nM PMA. Data represent mean ± SEM (n=4 mice), normalized to the unstimulated control condition. One-way ANOVA with Bonferroni post hoc test. NS, not significant; ***P<0.001.

We next measured binding of CD4^+^ T cells to soluble ICAM-1 (22), VCAM-1 (45), and MAdCAM-1 (16) to assess the activation of αLβ2, α4β1 and α4β7 integrins, respectively (Fig. 1B-D). Tln1-cR35E T cells exhibited a significant reduction in ICAM-1 (Fig. 1B) and MadCAM-1 (Fig. 1D) binding upon stimulation with phorbol myristate acetate (PMA). This defective binding was even more pronounced in Tln1-cR118E T cells. In sharp contrast, binding of mutant T cells to VCAM-1 was similar to control cells (Fig. 1C). Thus, these findings indicate that blockade of Rap1 binding to talin1 F0 or F1 subdomains partially inhibits activation of αLβ2 and α4β7 integrins, while preserving activation of α4β1 integrin.

### Blockade of Rap1 binding to both talin1 F0 and F1 subdomains markedly reduces integrin activation in CD4^+^ T cells

To assess the contribution of Rap1-talin1 interaction in T cells (Fig. 2A), we crossed *Tln1*^*WT/R35E,R118E*^*;CD4-Cre*^*+/-*^ mice (38) with the *Tln1*^*Flox/Flox*^ strain to obtain lymphocyte specific deletion of Tln1 flox allele in *Tln1*^*WT/Flox*^*;CD4-Cre*^*+/-*^ (control) and *Tln1*^*R35E,R118E/Flox*^*;CD4-Cre*^*+/-*^ (indicated as Tln1-cR35E,R118E) littermates for experiments. Lymphocyte content of talin1, RIAM and Rap1a/b (Supplementary Fig. 1C) and surface expression of αL, β2, α4, β1 and β7 integrins (Supplementary Fig. 1D) in Tln1-cR35E,R118E T cells were similar to wildtype T cells. Tln1-cR35E,R118E mice were viable, appeared healthy and exhibited normal counts of leukocytes in peripheral blood (Fig. 2B). Binding of PMA-stimulated Tln1-cR35E,R118E T cells to ICAM-1 and MAdCAM-1 was strongly reduced when compared to control T cells (Fig. 2C). Similarly, binding of Tln1-cR35E,R118E T cells to VCAM-1 was impaired (Fig. 2C), indicating that complete blockade of Rap1 binding to talin1 regulates α4β1 activation, in addition to αLβ2 and α4β7 integrins. Accordingly, adhesion of Tln1-cR35E,R118E T cells onto the three integrin ligands was impaired upon stimulation with either PMA or SDF1α under a flow condition (Fig. 2D). Since talin1 plays an important role in integrin-mediated trafficking of lymphocytes to lymph nodes (16, 46), we performed a competitive homing assay to measure the capacity of Tln1-cR35E,R118E T cells to migrate in vivo (Fig. 2E). Tln1-cR35E,R118E T cells manifested a reduced homing to both mesenteric and peripheral lymph nodes (Fig. 2F). Thus, disruption of the Rap1-talin1 interaction profoundly impairs integrin activation and function in CD4^+^ T cells.

**Figure 2.**
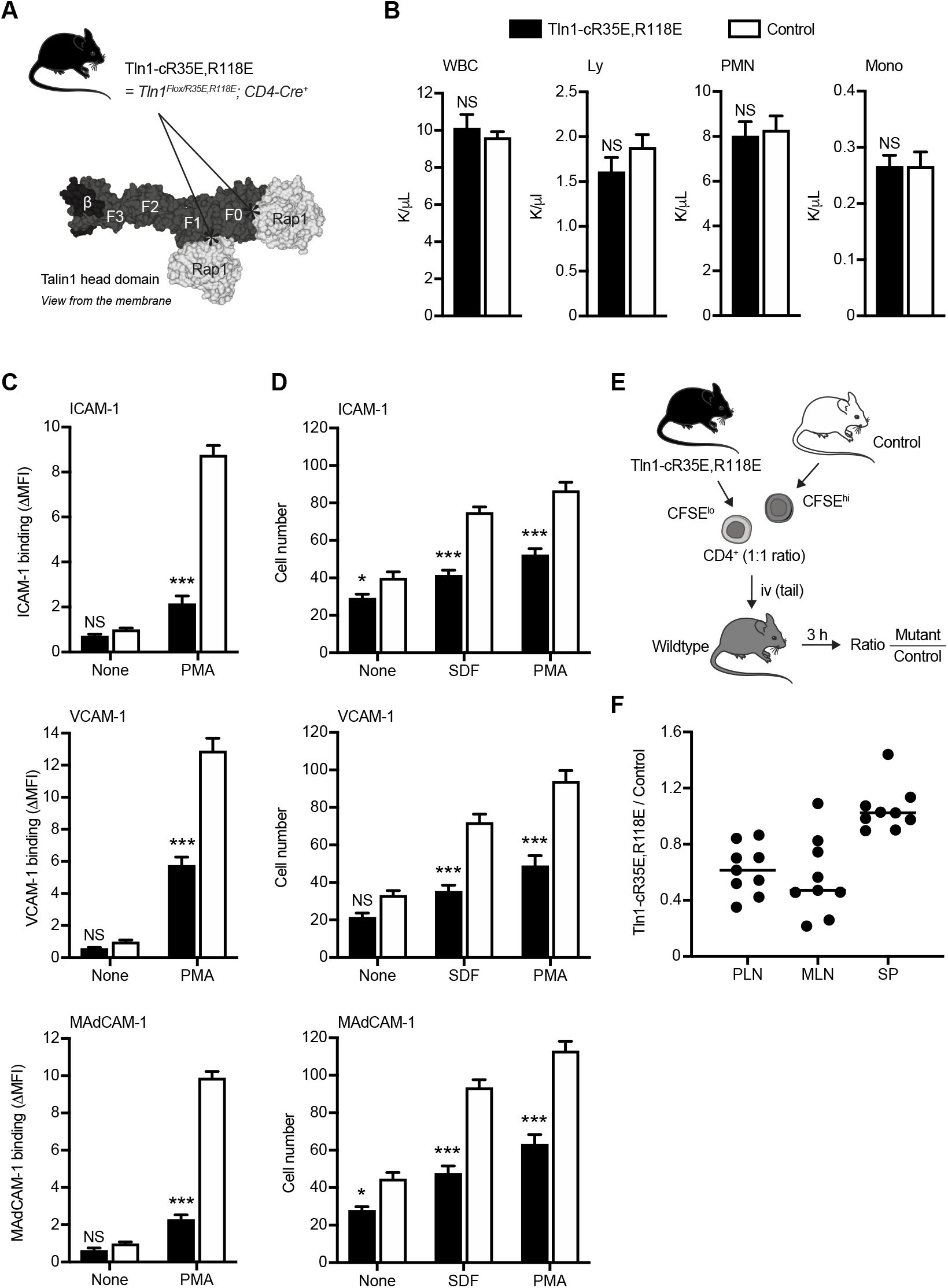
Blockade of Rap1 binding to both F0 and F1 domains in talin1 prevents integrin activation in CD4^+^ T cells. (A) Both R35E and R118E mutations in Talin1 F0 and F1 domains, respectively, prevents Rap1-talin1 interaction. (B) Peripheral blood cell counts of Tln1-cR35E,R118E mice. Values are mean ± SEM (n=6 mice). Two-tailed *t*-test; no significant differences were observed. WBC, white blood cells; PMN, polymorphonuclear neutrophils; Ly, lymphocytes; Mono, monocytes. (C) Binding of control or Tln1-cR35E,R118E mutant CD4^+^ T cells to soluble integrin ligands. Cells were stimulated with 100 nM PMA. Data represent mean ± SEM (n=10 mice), normalized to the unstimulated control condition. Two-way ANOVA with Bonferroni post hoc test. NS, not significant; ***P<0.001. (D) Adhesion of control or Tln1-cR35E,R118E mutant CD4^+^ T cells to integrin ligands under a wall shear stress of 2 dyn/cm^2^. Cells were stimulated with 1 μg/ml SDF1α or 100 nM PMA. The number of arrested adherent cells was plotted. Data are mean ± SEM (n=6 mice). Two-way ANOVA with Bonferroni post hoc test. NS, not significant; *P<0.05; ***P<0.001. (E-F) In vivo competitive homing of Tln1-cR35E,R118E mutant CD4^+^ T cells to different lymphoid tissues. CD4^+^ T cells were isolated from either control or Tln1-cR35E,R118E mice, differentially labeled and mixed prior to injection into C57BL/6 mice. The ratio of Tln1-cR35E,R118E mutant to control CD4^+^ T cells was determined by flow cytometry from lymph nodes 3 hrs after injection. Data represent mean ± SEM (n=9 mice).

### Both RIAM and Rap1 binding to talin1 operate in CD4^+^ T cells to regulate integrin activity

Because blockade of Rap1-talin1 interaction partially diminished integrin activation in Tln1-cR35E,R118E T cells, we reasoned that additional routes linking Rap1 to talin1 are involved in CD4^+^ T cells. RIAM is a member of the Mig-10/RIAM/LPD (MRL) family of adaptor proteins that participate to cell migration by controlling integrin activation and dynamics of the actin cytoskeleton (24, 25, 47, 48). We hypothesized that RIAM and talin1 together contributed to the capacity of Rap1 to promote integrin activation in CD4^+^ T cells (Fig. 3A). We crossed the Tln1-cR35E,R118E mice with the conditional *Apbb1ip* knockout strain, which is null for RIAM expression. *Tln1*^*R35E,R118E/Flox*^*;Apbb1ip*^*Flox/Flox*^*;CD4-Cre*^*+/-*^ mice (indicated as Tln1-cR35E,R118E; RIAM-cKO) were compared to *Tln1*^*WT/Flox*^*;Apbb1ip*^*WT/WT*^*;CD4-Cre*^*+/-*^ (control) littermates for experiments. These mutant mice were viable and apparently healthy, and their CD4^+^ T cells expressed a normal complement of talin1 and Rap1a/b (Supplementary Fig. 1E) or αL, β2, α4, β1 and β7 integrins (Supplementary Fig. 1F) when compared to control cells. Tln1-cR35E,R118E; RIAM-cKO T cells exhibited a substantial reduction in their capacity to home to peripheral and mesenteric lymph nodes (Fig. 3B). In agreement with previous reports (28, 32), we observed impaired binding of RIAM-deficient CD4^+^ T cells to ICAM-1 and MadCAM-1 in response to PMA stimulation, whereas defects in binding to VCAM-1 were minimal (Fig. 3C). The reduction in integrin ligand binding was greater in Tln1-cR35E,R118E T cells. These defects were even more dramatic in Tln1-cR35E,R118E; RIAM-cKO T cells in which binding to ICAM-1 and MadCAM-1 was completely abolished (Fig. 3C), thus mimicking the effect of Rap1a/b deletion. Thus, the combination of Rap1 binding to talin and to RIAM can account for the effect of Rap1 in CD4^+^ T cell integrin activation.

**Figure 3.**
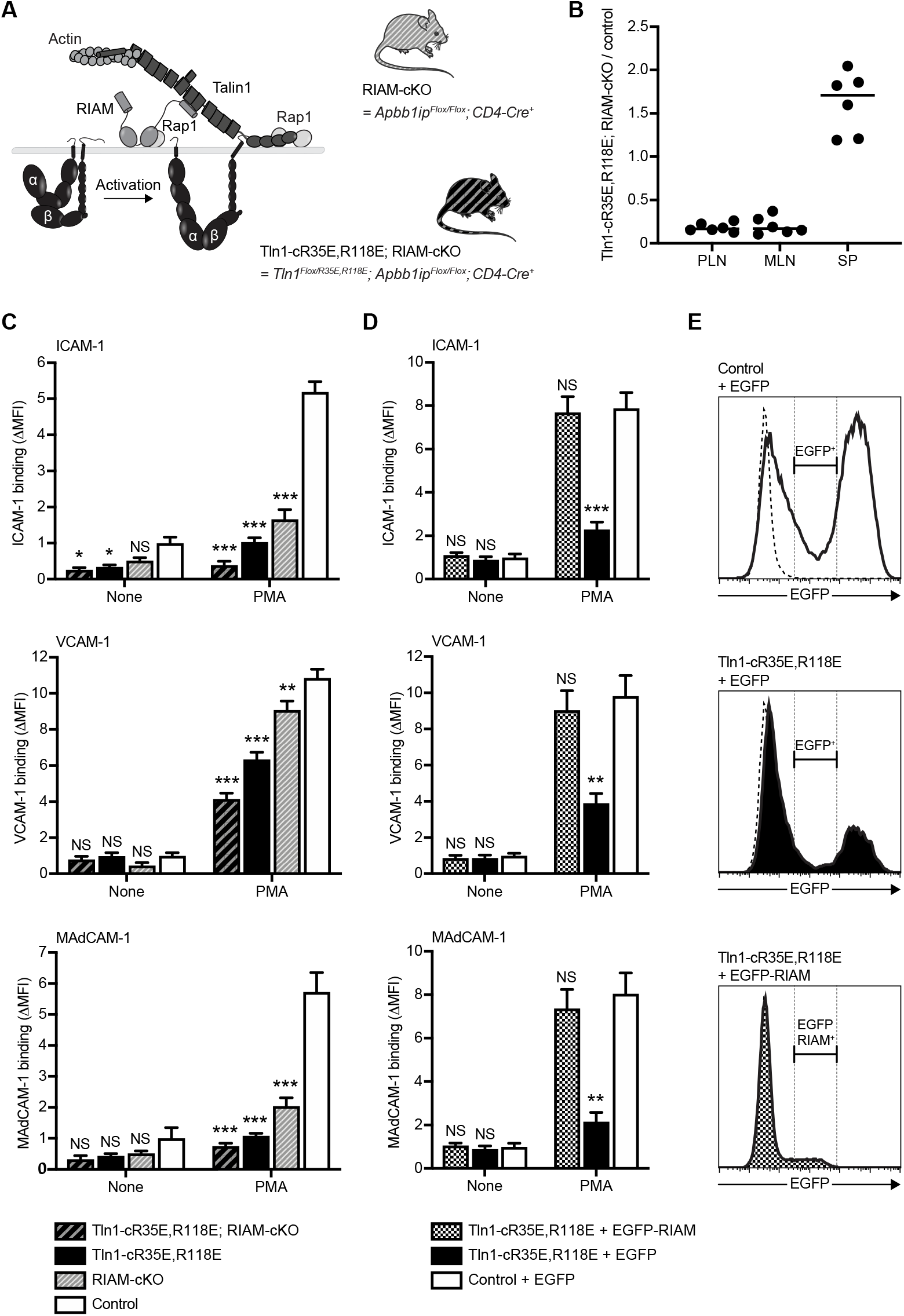
Rap1-talin1 direct interaction synergizes with Rap1-RIAM-talin1 to mediate integrin activation in CD4^+^ T cells. (A) RIAM function as a Rap1 effector that recruits talin1 to the plasma membrane, in parallel to Rap1 binding to talin1, to enable its interaction with the integrin β cytoplasmic tail. (B) In vivo competitive homing of Tln1-cR35E,R118E; RIAM-cKO CD4^+^ T cells to different lymphoid tissues. CD4^+^ T cells were isolated from either control or Tln1-cR35E,R118E; RIAM-cKO mice, differentially labeled and mixed prior to injection into C57BL/6 mice. The ratio of Tln1-cR35E,R118E; RIAM-cKO mutant to control CD4^+^ T cells was determined by flow cytometry from lymph nodes 3 hrs after injection. Data represent mean ± SEM (n=6 mice). (C) Binding of control, Tln1-cR35E,R118E or Tln1-cR35E,R118E; RIAM-cKO mutant CD4^+^ T cells to soluble integrin ligands. Cells were stimulated with 100 nM PMA. Data represent mean ± SEM (n=4 mice), normalized to the unstimulated control condition. Two-way ANOVA with Bonferroni post hoc test. NS, not significant; *P<0.05; **P<0.01; ***P<0.001. (D-E) CD4^+^ T cells isolated from control or Tln1-cR35E,R118E mice were transduced with retroviruses encoding either EGFP-tagged RIAM or EGFP alone. Integrin activation in EGFP^+^ cells was analyzed by flow cytometry. (D) The level of EGFP expression was determined at 3 days after transduction. (E) Binding of transduced CD4^+^ T cells to soluble integrin ligands. Cells were stimulated with 100 nM PMA. Data represent mean ± SEM (n=4 mice). Two-way ANOVA with Bonferroni post hoc test. NS, not significant; **P<0.01; ***P<0.001.

We reasoned that if talin1 or RIAM could connect Rap1 to integrins (Fig. 3A), then RIAM might be able to bridge Rap1 to talin1(R35E,R118E) thereby compensating for the loss of direct Rap1 binding to talin1(R35E,R118E). We therefore tested whether increasing RIAM expression in Tln1-cR35E,R118E CD4^+^ T cells can restore the activation defects of αLβ2, α4β1 and α4β7 integrins. We transduced proliferating Tln1-cR35E,R118E CD4^+^ T cells with MSCV retroviruses encoding either RIAM-IRES-EGFP or EGFP alone. Overexpression of RIAM in Tln1-cR35E,R118E T cells fully restored activation of αLβ2, α4β1 and α4β7 integrins (Fig. 3D-E). These data demonstrate that both Rap1-RIAM axis and Rap1 binding to talin1 contribute to integrin activation in CD4^+^ T cells, with Rap1-talin1 interaction making a greater contribution and that RIAM can compensate for the defect in integrin action in Tln1-cR35E,R118E CD4^+^ T cells.

### Rap1-talin1 interaction participates to integrin activation in Treg cells

We previously reported distinct integrin activation pathways operate in effector and Treg cells (32). We crossed *Tln1*^*WT/R35E,R118E*^ mice with *Foxp3*^*YFP-Cre*^ mice to generate *Tln1*^*WT/R35E,R118E*^*;Foxp3*^*YFP-Cre*^ mice in which the Cre recombinase is selectively expressed in Treg cells. We next crossed *Tln1*^*WT/R35E,R118E*^*;Foxp3*^*YFP-Cre*^ mice with the *Tln1*^*Flox/Flox*^ strain to obtain Treg cell specific deletion of Tln1 flox allele in *Tln1*^*WT/Flox*^*;Foxp3*^*YFP-Cre*^ (control) and *Tln1*^*R35E,R118E/Flox*^*;Foxp3*^*YFP-Cre*^ (indicated as Tln1rR35E,R118E) littermates for experiments (Fig. 4A). Expression of Talin1(R35E,R118E), RIAM and Rap1a/b was unaffected in Tln1-rR35E,R118E Treg cells when compared to *Tln1*^*WT/Flox*^; *Foxp3*^*YFP-Cre*^ control Treg cells (Supplementary Fig. 2A), and surface levels of αL, β2, α4, β1 and β7 integrins was intact (Supplementary Fig. 2B). Tln1-rR35E,R118E mice manifested a significant leukocytosis (Fig. 4B), which mirrors a previous observation that loss of talin1 function in Treg cells causes systemic inflammation (20, 21). Binding of ICAM-1 and MAdCAM-1 to PMA stimulated Tln1-rR35E,R118E Treg cells was diminished by 50%, whereas binding of VCAM-1 was similar to control Treg cells (Fig. 4C). Consistent with impaired integrin activation, Tln1-rR35E,R118E Treg cells homed less efficiently to peripheral and mesenteric lymph nodes (Fig. 4D). Because both Rap1 and talin1 are required to exert suppressive function in Treg cells, we next performed an in vitro suppression assay (Fig. 4E). We observed that Tln1-rR35E,R118E Treg cells exhibited reduce capacity to suppress proliferation of conventional T cells (Fig. 4F). Together, our findings reveal that Rap1-talin1 interaction in Treg cells contributes to the activation and associated functions of αLβ2 and α4β7 integrins, whereas, unlike in CD4^+^ T cells, α4β1 activity is preserved.

**Figure 4.**
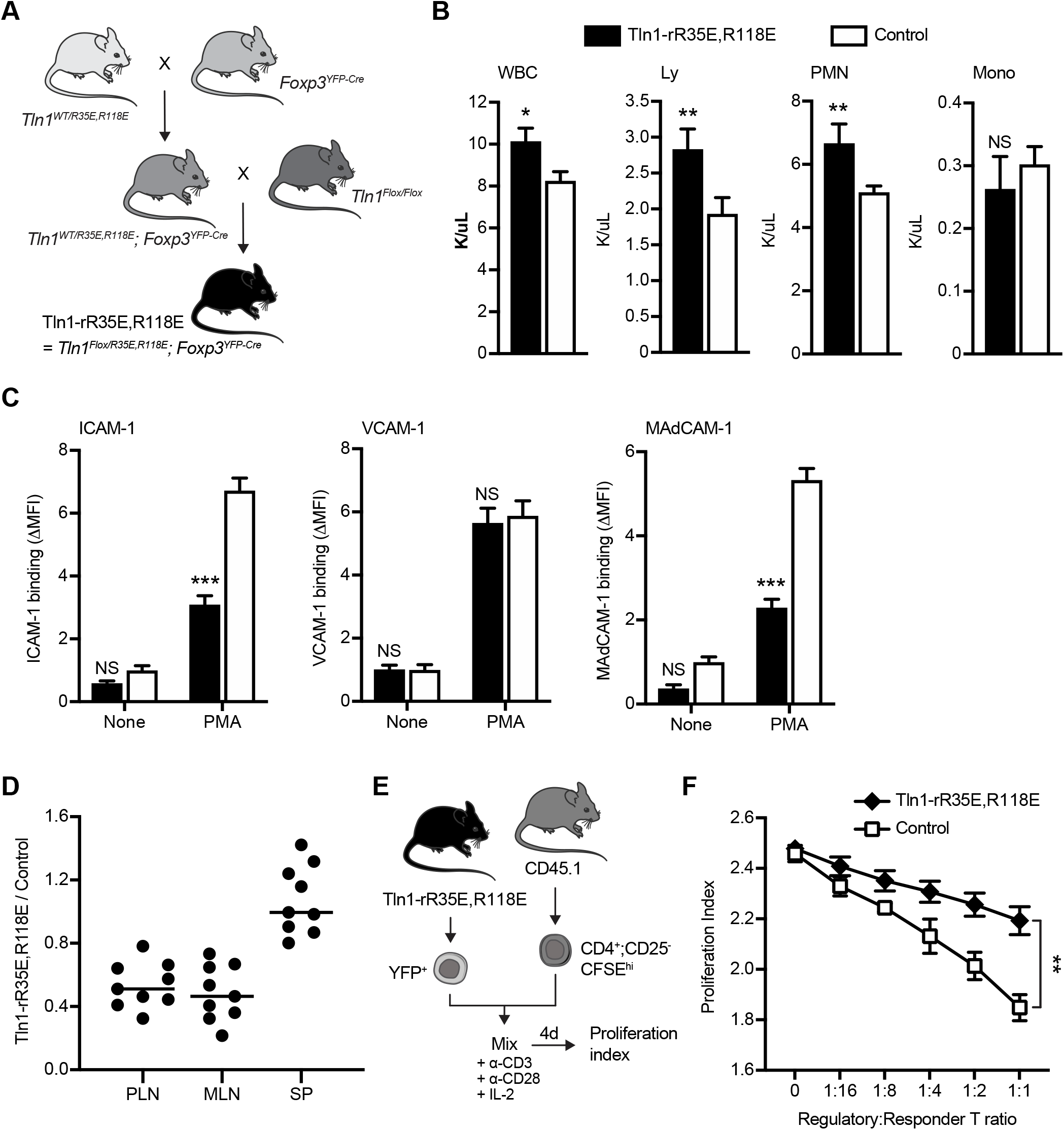
Rap1 binding to talin1 participates to activate integrins in Treg cells. (A) Mouse breeding strategy to selectively restrict the expression of talin1(R35E,R118E) in Treg cells. (B) Peripheral blood cell counts of Tln1-rR35E,R118E mice. Values are mean ± SEM (n=6 mice). Two-tailed *t*-test; NS, not significant; *P<0.05; **P<0.01. WBC, white blood cells; PMN, polymorphonuclear neutrophils; Ly, lymphocytes; Mono, monocytes. (C) Binding of control or Tln1-rR35E,R118E mutant Treg cells to soluble integrin ligands. Cells were stimulated with 100 nM PMA. Data represent mean ± SEM (n=8 mice), normalized to the unstimulated control condition. Two-way ANOVA with Bonferroni post hoc test. NS, not significant; ***P<0.001. (D) In vivo competitive homing of Tln1-rR35E,R118E mutant Treg cells to different lymphoid tissues. Treg cells were isolated from either control or Tln1-rR35E,R118E mice, differentially labeled and mixed prior to injection into C57BL/6 mice. The ratio of Tln1-rR35E,R118E mutant to control Treg cells was determined by flow cytometry from lymph nodes 3 hrs after injection. Data represent mean ± SEM (n=9 mice). (E-F) Treg cell suppressive activity. Treg cells isolated from CD45.2 congenic control or Tln1-rR35E,R118E mice were mixed with Responder T cells at the indicated Treg/Responder cell ratios. Responder cells were CFSE-labelled CD4^+^CD25^-^ naive T cells isolated from CD45.1 congenic C57BL/6 mice and activated by anti-CD3 (5 μg/ml), anti-CD28 (5 μg/ml) and IL-2. CFSE^+^ populations gated on CD45.1^+^ cells were analyzed by flow cytometry at day4 to determine the proliferation index using FlowJo software (n=3 mice).

### RIAM and LPD complement Rap1-talin1 interaction in promoting integrin activation and function in Treg cells

RIAM deficiency has a minor effect on Treg cell integrin activation and function (32). To ask whether Rap1 binding to talin1 supports integrin activation in RIAM-null Treg cells, we generated *Tln1*^*R35E,R118E/Flox*^*;Apbb1ip*^*Flox/Flox*^*;Foxp3*^*YFP-Cre*^ mice (indicated as Tln1-rR35E,R118E; RIAM-rKO) and *Tln1*^*WT/Flox*^*;Apbb1ip*^*WT/WT*^*;Foxp3*^*YFP-Cre*^ (control) littermates for experiments. Tln1-rR35E,R118E; RIAM-rKO Treg cells express a similar level of talin1 as in *Tln1*^*WT/Flox*^; *Foxp3*^*YFP-Cre*^ Treg cells and equal content of Rap1a/b (Supplementary Fig. 2C) or surface αL, β2, α4, β1 and β7 integrins (Supplementary Fig. 2D) when compared to control Treg cells. Tln1-rR35E,R118E; RIAM-rKO mice were viable; however, they exhibited a reduced body size when compared to control mice (Fig. 5A) and manifested larger lymph nodes and spleen but smaller thymus (Fig. 5B), suggesting that these mice experienced immune dysregulation. Treg cells isolated from Tln1-rR35E,R118E; RIAM-rKO mice showed reduced binding to ICAM-1 and MAdCAM-1 after PMA stimulation to a similar extent to that observed in Tln1-R35E,R118E Treg cells (Fig. 5C). However, Tln1-rR35E,R118E; RIAM-rKO Treg cells also exhibited a 50% reduction in binding to VCAM-1, explaining why loss of RIAM could impact the function of Tln1-rR35E,R118E Treg cells (Fig. 5C). In contrast, *Apbb1ip*^*Flox/Flox*^*;Foxp3*^*YFP-Cre*^ mice (indicated as RIAM-rKO) displayed normal binding to soluble integrin ligands (Fig. 5C) as previously described (32). Moreover, Treg cells lacking Rap1-talin1 interaction and RIAM were not able to suppress the proliferation of conventional T cells (Fig. 5D) and homed less to peripheral and mesenteric lymph nodes (Fig. 5E). These data suggest that although RIAM is dispensable for integrin activation in Treg cells, it complements the direct binding of talin1 to Rap1 in supporting Treg functions.

**Figure 5.**
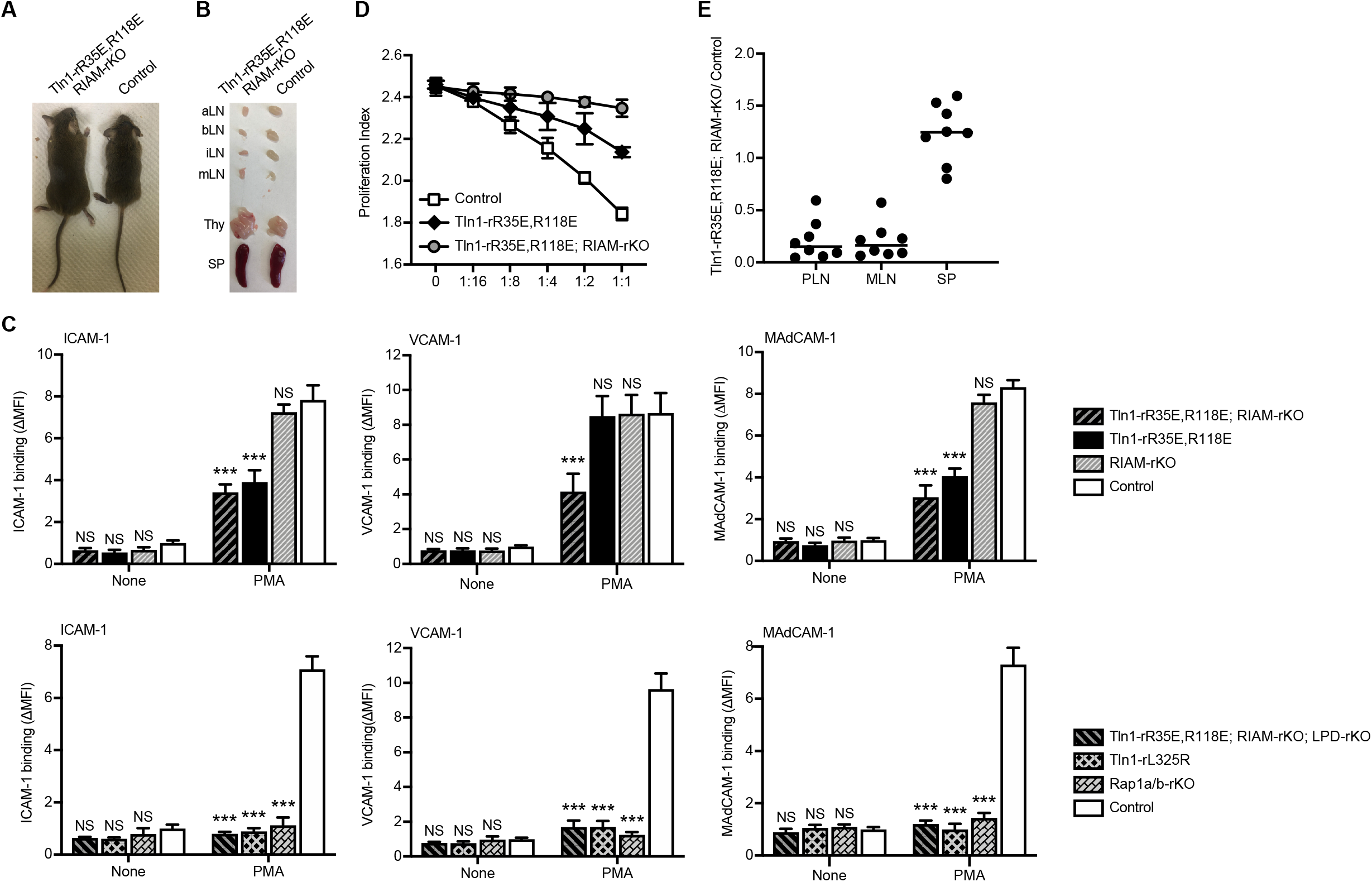
Both LPD and Rap1 binding to talin1 contribute to activate integrins in Treg cells. (A-B) Representative morphology (A) and lymphoid tissues (B) of Tln1-rR35E,R118E; RIAM-rKO mice. aLN, aortic lymph nodes; bLN, brachial lymph nodes; iLN, inguinal lymph nodes; mLN, mesenteric lymph nodes; Thy, thymus; SP, spleen. (C) Binding of control, RIAM-rKO, Tln1-rR35E,R118E or Tln1-rR35E,R118E; RIAM-rKO mutant Treg cells to soluble integrin ligands. Cells were stimulated with 100 nM PMA. Data represent mean ± SEM (n=4 mice), normalized to the unstimulated control condition. Two-way ANOVA with Bonferroni post hoc test. NS, not significant; ***P<0.001. (D) Treg cell suppressive activity. Treg cells isolated from CD45.2 congenic control or Tln1-rR35E,R118E; RIAM-rKO mice were mixed with Responder T cells at the indicated Treg/Responder cell ratios. Responder cells were CFSE-labelled CD4^+^CD25^-^ naive T cells isolated from CD45.1 congenic C57BL/6 mice and activated by anti-CD3 (5 μg/ml), anti-CD28 (5 μg/ml) and IL-2. CFSE^+^ populations gated on CD45.1^+^ cells were analyzed by flow cytometry at day4 to determine the proliferation index using FlowJo software (n=4 mice). (E) In vivo competitive homing of Tln1-rR35E,R118E; RIAM-rKO Treg cells to different lymphoid tissues. Treg cells were isolated from either control or Tln1-rR35E,R118E; RIAM-rKO mice, differentially labeled and mixed prior to injection into C57BL/6 mice. The ratio of Tln1-rR35E,R118E; RIAM-rKO mutant to control Treg cells was determined by flow cytometry from lymph nodes 3 hrs after injection. Data represent mean ± SEM (n=8 mice). (F) Binding of control, Tln1-rL325R or Tln1-rR35E,R118E; RIAM-rKO; LPD-rKO mutant Treg cells to soluble integrin ligands. Cells were stimulated with 100 nM PMA. Data represent mean ± SEM (n=4 mice). Two-way ANOVA with Bonferroni post hoc test. NS, not significant; ***P<0.001.

The foregoing results indicate that direct binding of Rap1 to talin1 is important in Treg and Tconv integrin activation; however, the MRL proteins, RIAM and LPD, can also subserve this function. Furthermore, in Tconv, RIAM and talin1 together can completely account for the capacity of Rap1 to support integrin activation. Because of LPD’s importance in Treg cells, we generated *Tln1*^*R35E,R118E/Flox*^*;Apbb1ip*^*Flox/Flox*^*;Raph1*^*Flox/Flox*^*;Foxp3*^*YFP-Cre*^ mice (indicated as Tln1-rR35E,R118E; RIAM-rKO; LPD-rKO) and *Tln1*^*WT/Flox*^*;Apbb1ip*^*WT/WT*^*;Raph1*^*WT/WT*^*;Foxp3*^*YFP-Cre*^ (control) littermates for experiments. Tln1-rR35E,R118E; RIAM-rKO; LPD-rKO Treg cells exhibited a complete loss of activation of α4β1, αLβ2 and α4β7 integrins (Fig. 5F). The defects were similar to that observed in Treg cells lacking Rap1a/b (indicated as Rap1a/b-rKO), or isolated from *Tln1*^*L325R/Flox*^*;Foxp3*^*YFP-Cre*^ mice (indicated as Tln1-rL325R) harboring a mutation that blocks talin1’s capacity to activate integrins (40, 49). Taken together, these results establish that the combination of direct binding of Rap1 to talin1 with LPD and RIAM can account for the role of Rap1 in integrin function in Treg cells.

## Discussion

Rap1 is a major convergence point of the lymphocyte signaling pathways that result in talin1 binding to the integrin β cytoplasmic tail and subsequent integrin activation to shape a successful immune response. Although the contribution of RIAM, which is a Rap1 effector, to trigger talin1-dependent activation of T cell integrins is well documented (28, 29, 31, 32, 48), the nature of the connection between Rap1 and talin1 in lymphocytes remains incompletely characterized. We recently identified two Rap1 binding sites in talin1 F0 and F1 subdomains and generated mice bearing the two point mutations R35E and R118E in talin1 F0 and F1 subdomains (35, 37), respectively, which block binding to Rap1 (38). Here, we assessed the contribution of the Rap1-talin1 interaction to trigger integrin activation in T cells and report a thorough analysis of the dialogue between Rap1 and talin1 in conventional CD4^+^ lymphocytes or Treg cells. Disruption of Rap1 binding to talin1 causes a significant, but moderate, defect in αLβ2, α4β1 and α4β7 integrin activation in CD4^+^ T cells. RIAM null T cells expressing the talin1(R35E,R118E) mutant manifest further impaired integrin activation. Inversely, overexpression of RIAM restored the capacity of the talin1(R35E,R118E) mutant to mediate integrin activation. Similarly, Rap1-talin1 interaction participates, in parallel with LPD, but not RIAM, to αLβ2 and α4β7 integrin function in Treg cells. These data reveal the contribution of multiple routes that connect Rap1 to talin1 in Treg and CD4^+^ T cells.

Blockade of Rap1 binding to talin1 F0 or F1 subdomains partially inhibits activation of αLβ2 and α4β7 integrins in T cells, while the loss of both Rap1 binding sites further impairs integrin activation. This finding is consistent with our previous observation that disabling both Rap1 binding sites in F0 and F1 subdomains had a greater effect than hindering F0 or F1 alone (38). Likewise, Bromberger and colleagues reported that a talin1 knock-in mouse strain (Tln1^3mut^), which carries mutations in talin1 F0 subdomain to block Rap1 binding, manifest mild defects in platelet integrin functions (36), similar to mice expressing the talin1(R35E) mutant (35). Furthermore, blockade of the Rap1-talin1 F0 interaction led to normal blood cell counts in the peripheral blood of talin1(R35E) expressing mice (35) or Tln1^3mut^ mice (36). In addition, neutrophils from the Tln1^3mut^ mouse have a mild adhesion defect and manifest a partial reduction in blood extravasation (36). However, talin1(R118E) expressing mice with loss of the Rap1 binding site in talin1 F1 alone exhibit a significant leukocytosis that affects both neutrophils and lymphocytes (38). In sum, these data suggest that the Rap1-talin1 F1 interaction has a greater functional impact than the Rap1-talin1 F0 interaction; however, both Rap1 binding sites in talin1 F0 and F1 cooperate to potentiate integrin function in T cells.

Loss of the Rap1-talin1 interaction binding did not affect the activation of integrin α4β1 in Treg cells and led to a partial defect in CD4^+^ T cells, unlike αLβ2 and α4β7. Previous studies demonstrated that deletion of RIAM in T cells partially suppressed αLβ2 and α4β7 integrin activation but did not block activation of α4β1 (28, 29, 32). In sharp contrast, deletion of either both Rap1a and Rap1b isoforms or talin1 profoundly suppressed activation of α4β1 in addition to the other two integrins (28, 29). The loss of homing of these mutant T cells to peripheral lymphoid organs was correlated with a loss of integrin activation. These data indicate that another Rap1 effector is involved in β1 integrin activation. RapL is a Rap1 effector that regulates the activity of αLβ2 integrin through binding to the αL cytoplasmic tail (50, 51). It appears to regulate kindlin-3 rather than talin1 (52). Previous studies have shown that RAPL-deficient T lymphocytes had impaired adhesion to VCAM-1 upon stimulation with chemokine ligand 21 (CCL21) despite having a normal surface expression of VLA-4. In addition, RAPL-deficient T lymphocytes also showed defective abilities of homing to peripheral lymph nodes and trafficking to spleens which require activation of αLβ2 and α4β1 (51). RAPL deficiency also inhibited cell adhesion to VCAM-1 in the human T cell line Molt-4 (53). Taken together, these findings raise the possibility that RAPL contributes to the activation of integrin α4β1 while compensating for the lack of RIAM and Rap1-talin1 interaction in T cells. Additional studies are required to address this point and establish the relative importance of each talin1-dependent signaling modules with respect to the RapL pathway.

While the Rap1-RIAM axis is partially redundant to the Rap1-talin1 interaction in CD4^+^ T cells, RIAM is dispensable in Treg cells. Instead, LPD functions in parallel to the Rap1-talin1 interaction in Treg cells. In conclusion, our findings reveal that although both Rap1 and talin1 are essential for blood cell integrin functions, their connection differ in a cell-type specific manner. These differences embody potential avenues to manipulate the functions and trafficking of particular lymphocyte subsets for fine tuning the immune response.

## Acknowledgements

The authors thank Mark Ginsberg for helpful discussions and support, and critical reading of the manuscript. This work was supported by the American Heart Association Career Development Award 18CDA34110228 (F.L.), the European Union’s Horizon 2020 research and innovation programme – Marie Skłodowska-Curie grant No 841428 (F.L.), National Institutes of Health – National Heart, Lung and Blood Institute grant KO1 HL133530, P01HL151433-01 (M.A.L.R.), HL139947 (M.H.G.) and HL145454 (Z.C.F.).

## Footnotes

### Contribution

H.S. supervised the research; H.S. and F.L. conceived the study, designed experiments, interpreted data, and wrote the manuscript; F.L., H.S., B.Y.T, Q.Y.D., and W.W.Q. performed and analyzed experiments; Z.C.F., M.A.L.R. and A.R.G provided vital reagents and critical expertise.

## Supplemental Information

**Supplementary Figure 1.**
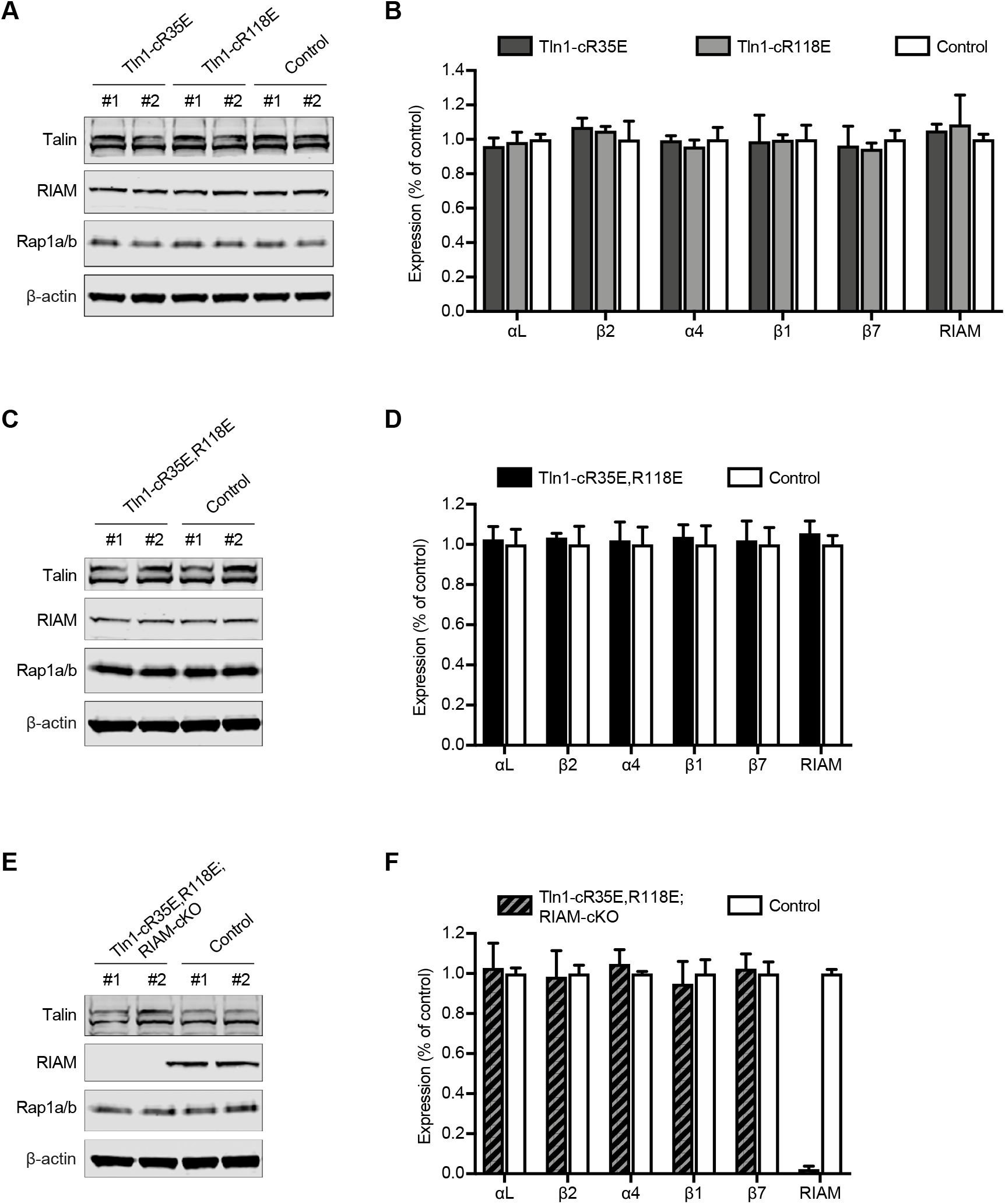
Intact expression of talin1(R35E,R118E) mutant in CD4^+^ T cells. Expression of talin1, RIAM and Rap1a/b in Tln1-cR35E or Tln1-cR118E **(A)**, Tln1-cR35E-R118E **(C)** and Tln1-R35E,R118E; RIAM-cKO **(E)** mutant CD4^+^ T cells was assayed by Western blotting. Results are representative of 3 independent experiments, n=2 mice each time. Surface expression of integrins in Tln1-cR35E or Tln1-cR118E **(B)**, Tln1-cR35E-R118E **(D)** and Tln1-R35E,R118E; RIAM-cKO **(F)** mutant CD4^+^ T cells was measured by flow cytometry. Bar graphs represent MFI +/-SEM (n=3 mice). Two-tailed t-test; no significant differences were observed.

**Supplementary Figure 2.**
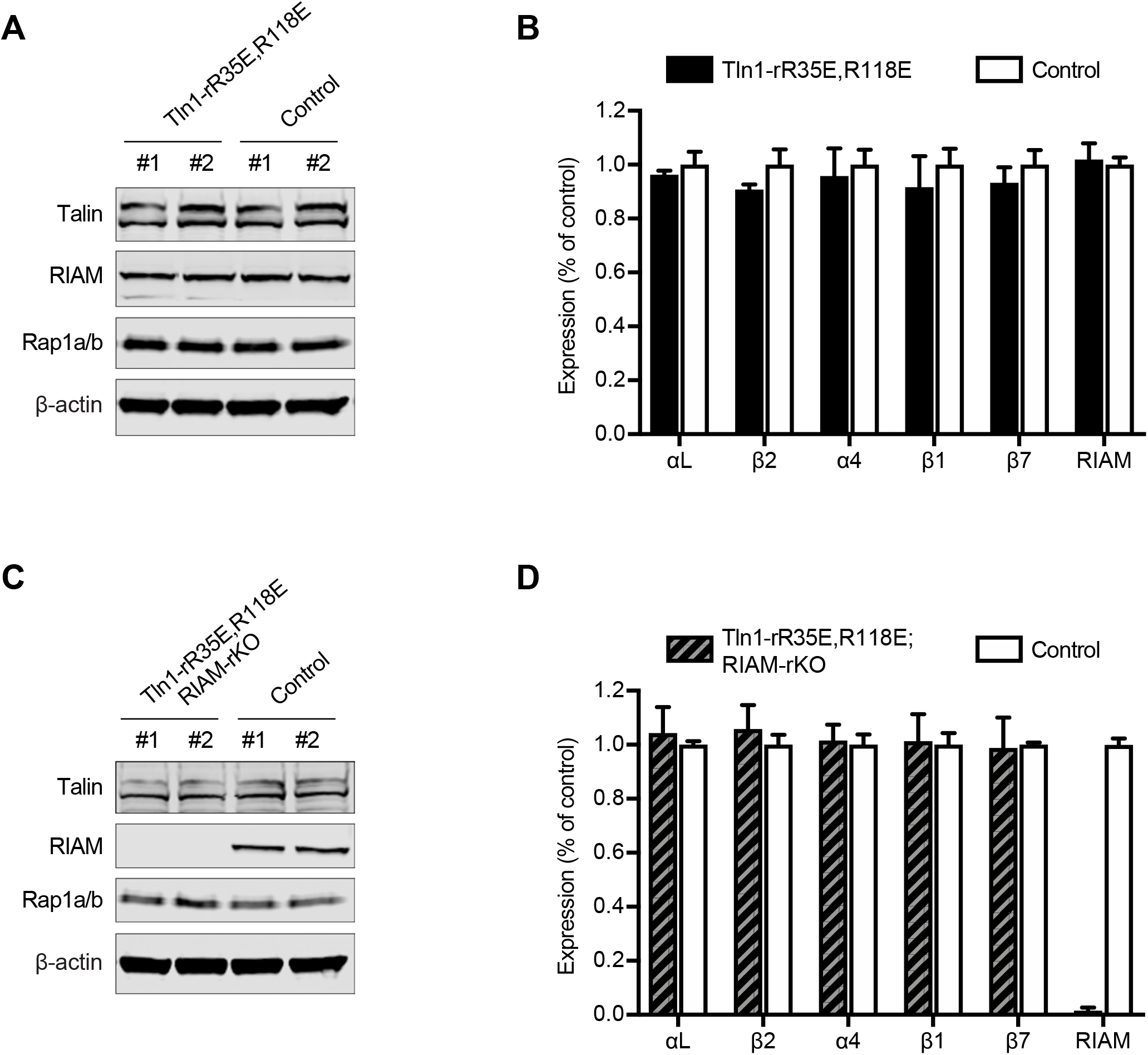
Intact expression of talin1(R35E,R118E) mutant in Treg cells. **(A**,**C)** Expression of talin1, RIAM and Rap1a/b in Tln1-rR35E,R118E **(A)** or Tln1-R35E,R118E; RIAM-rKO **(C)** mutant Treg cells was assayed by Western blotting. Results are representative of 3 independent experiments, n=2 mice each time. **(B**,**D)** Surface expression of integrins in Tln1-rR35E,R118E **(B)** or Tln1-R35E,R118E; RIAM-rKO **(D)** mutant CD4^+^ T cells was measured by flow cytometry. Bar graphs represent MFI +/-SEM (n=3 mice). Two-tailed t-test; no significant differences were observed.

